# Quantum implementation of multi-pattern string matching for k-mer detection

**DOI:** 10.64898/2025.12.21.695808

**Authors:** Christos Papalitsas, Ioannis Mouratidis, Michail Patsakis, Evangelos Stogiannos, Ilias Georgakopoulos-Soares, Grigorios Koulouras

**Affiliations:** Division of Pharmacology and Toxicology, College of Pharmacy, The University of Texas at Austin, Dell Paediatric Research Institute, Austin, TX, USA

## Abstract

**Motivation:** The exponential growth of publicly available genomic data has created unprecedented opportunities for sequence-based discovery. Locating specific k-mers is fundamental to diverse applications, including metagenomic classification, pathogen and cancer detection, and variant calling yet efficient identification of multiple k-mer patterns across large sequencing data and massive databases remains a significant computational challenge.

**Method:** We implement two quantum algorithms for DNA multi pattern string matching for k-mer detection based on Grover’s amplitude amplification with quantum random access memory (QRAM). The first algorithm uses an enumerate-m oracle that sequentially checks a loaded text substring against all m patterns achieving O(√S) query complexity for S text positions but requiring O(m·L) work per oracle call. The second algorithm employs nested Grover search with an outer loop over text positions and an inner loop over pattern space, reducing oracle complexity to O(L) while performing O(√S · √m) in total.

**Results:** We present two quantum implementations of multi-pattern string matching tailored for k-mer detection. Leveraging quantum parallelism and Grover-inspired search primitives, our methods accelerate dictionary-based pattern matching, particularly in contexts involving large sequences, such as genomic data, and extensive pattern sets.

**Conclusions:** While implementation challenges such as QRAM overhead remain, this study demonstrates both the promise and current limitations of quantum-enhanced string matching, establishing a foundational step toward quantum readiness in bioinformatics.

**Availability and implementation:** To maximize accessibility and practical use, we provide our methodology at: https://github.com/Georgakopoulos-Soares-lab/quantum-multi-motif-finder

## 1 Introduction

The rapid expansion of genomic sequencing technologies has generated vast resources of biological sequence data, creating both opportunities and computational challenges for modern bioinformatics. A fundamental task across diverse applications is the identification of specific k-mer sequences, short DNA or RNA subsequences of length k, within large sequencing datasets and massive databases. Variant detection relies on identifying k-mers that span or flank genomic variation to distinguish these altered regions from reference genomes (1, 2). In pathogen identification, detecting characteristic k-mers enables rapid species classification and strain typing, critical for outbreak response and clinical diagnostics (3, 4). Specifically, in metagenomic studies, where environmental samples contain nucleic acids from diverse organisms, the presence or absence of taxon-specific k-mers allows community composition analysis without prior cultivation or assembly. In cancer detection, cancer-specific k-mers can be leveraged for early detection (5, 6). As sequencing databases continue to grow, with repositories such as the NCBI Sequence Read Archive now exceeding tens of petabases, the computational burden of k-mer searches can become a bottleneck limiting analysis and large-scale comparative studies (7).

Quantum computing harnesses the principles of superposition, entanglement, and interference to perform computations that, in certain contexts, may surpass the capabilities of classical computing (8). Recent advances in quantum computing are unlocking promising opportunities to optimize bioinformatics algorithms, including those based on k-mer analysis (9, 10). Quantum algorithms offer promising speedups for problems such as string matching, an essential component of many sequence analysis pipelines that traditionally scale poorly with increasing dataset size (11). Prior research has demonstrated that quantum-enhanced pattern matching could significantly accelerate the identification of motifs across large genomic datasets. Approaches like QOMIC demonstrate how quantum circuits can efficiently solve motif detection problems, while quantum motif clustering methods apply Grover’s search and approximate counting to enhance motif-based graph analysis. These advances hold promise for improving the scalability and efficiency of downstream tasks such as metagenomic classification and transcriptome profiling (12, 13). For instance, a notable advancement in this direction is the recent breakthrough by Google’s Quantum AI team, who successfully demonstrated the Quantum Echoes algorithm on their Willow processor. Inspired by sonar-like interference patterns, this algorithm achieved a quantum advantage in simulating molecular structures, performing computations up to 13,000 times faster than classical supercomputers (14). Although Quantum Echoes has not yet been applied directly to bioinformatics, its core principle of using interference to extract meaningful patterns from complex quantum systems holds great potential for biological data analysis, particularly in scenarios involving noisy, high-dimensional datasets where signal extraction remains a persistent challenge.

The Aho-Corasick algorithm, developed by Alfred V. Aho and Margaret J. Corasick in 1975, is a seminal contribution to the field of classical string matching, particularly for applications requiring the simultaneous search of multiple patterns (15). Unlike traditional single-pattern algorithms, Aho-Corasick constructs a finite state machine that enables the detection of all occurrences of multiple patterns in a single linear pass through the input text. This makes exceptionally efficient for large-scale text processing tasks. However, as genomic datasets continue to grow exponentially, with modern sequencing projects generating petabytes of data, even the linear time complexity of classical algorithms like Aho-Corasick can become a computational bottleneck, motivating the exploration of quantum computing approaches that promise theoretical speedups through fundamentally different computational paradigms.

In this study we developed and benchmarked two distinct quantum algorithms for k-mer mapping in genomic sequences: the *Enumerate-m* algorithm and the *Nested Grover* algorithm. Both approaches leverage quantum superposition and amplitude amplification to efficiently search for multiple k-mer patterns simultaneously within a target DNA sequence. The fundamental challenge addressed in this work is how to exploit quantum parallelism to accelerate the traditionally computationally intensive task of finding all occurrences of multiple short DNA sequences (k-mers) within larger genomic texts. The implementations utilize quantum random access memory (QRAM) for sequence data storage and retrieval, combined with Grover’s algorithm for amplitude amplification. The two proposed algorithms differ significantly in their approach to pattern handling. The *Enumerate-m* algorithm stores patterns directly in the quantum circuit through hardcoded oracle operations, while the *Nested Grover* algorithm employs QRAM for both sequence and pattern storage. While classical approaches to multi-pattern matching scale linearly with both the number of patterns and the length of the target sequence, these quantum algorithms offer the potential for quadratic speedups through quantum superposition and amplitude amplification techniques.

## 2 Materials and methods

### 2.1. Harnessing quantum computing for genomic pattern matching

The identification of short DNA sequences, or k-mers, is a fundamental task in genomic analysis, supporting applications such as locating transcription factor binding sites, detecting repetitive elements, and developing diagnostic markers. Classical multiple pattern matching algorithms, such as Aho-Corasick, address this challenge efficiently. However, their runtime scales linearly with the length of the genomic sequence (n), which can become a limiting factor when processing large genomes or extensive k-mer libraries. As genomic datasets continue to grow in size and complexity, the need for more scalable computational paradigms becomes increasingly pressing.

Quantum computing enables parallel processing across multiple states by harnessing superposition, entanglement, and interference. In the context of genomic pattern search, this enables the simultaneous evaluation of all possible starting positions within a sequence, reducing the search time to scale with (√n). Such a quadratic speedup over classical methods holds significant promise for accelerating large-scale motif discovery, particularly when integrated with quantum memory architectures that allow efficient access to sequence data. This potential makes quantum algorithms a compelling avenue for advancing high-throughput genomic analysis (10).

Several quantum algorithms could, in theory, address the multiple pattern matching problem, but their suitability varies. The Quantum Fourier Transform (QFT), demonstrated experimentally by Weinstein et al. (16), is highly effective for period-finding and phase estimation, yet offers no direct advantage for arbitrary text matching without significant reformulation. Quantum walks, as reviewed by Venegas-Andraca (17), can traverse graph structures like suffix trees or automata, aligning conceptually with Aho-Corasick. However, their implementation demands complex state preparation and oracle design, making them less practical for near-term quantum devices.

Grover’s algorithm, by contrast, provides a clear, well-characterized quadratic speedup for unstructured search (15). Pattern matching can be naturally framed as searching among (n - L + 1) possible starting positions, enabling a simple oracle that flags matches. Its compatibility with QRAM allows efficient substring retrieval during the search, combining theoretical clarity with implementation feasibility. This makes Grover’s approach a pragmatic and scalable choice over more structurally complex alternatives.

### 2.2 Quantum random access memory (QRAM) and bucket-brigade architecture

QRAM is the quantum analogue of classical RAM. It’s designed to store data in superposition, meaning multiple memory addresses can be accessed simultaneously. Instead of binary addresses (0s and 1s), QRAM uses quantum addresses (i.e., superpositions of states) (18). This allows parallel access to multiple memory cells at once. A query input of ∑_i_ α_i_ |_i_⟩ |0⟩ returns the output ∑_i_ α_i_ |_i_⟩ |d_i_⟩, where d_i_ is the classical data at memory address i (19, 20). QRAM is still in its early stages, but ongoing research highlights its promise as a key enabler for quantum computing. Proposed architectures such as tree-based QRAM and circuit-based approaches aim to allow efficient quantum data access in superposition (18). While large-scale, fault-tolerant QRAM is not yet available for public or commercial use, small experimental demonstrations and theoretical models continue to advance, addressing challenges of error rates, hardware complexity, and scalability (21, 22). Current quantum systems rely on classical memory and limited quantum registers, yet QRAM research suggests future integration could unlock powerful applications in quantum machine learning, search, and simulation. To this end, a central premise of our methodology is the assumption of an efficient QRAM.

The bucket-brigade architecture represents a pivotal design for making large-scale QRAM physically realizable by significantly reducing the number of active components required per query. In this framework, memory is arranged as a binary tree of routing elements, each storing partial address information (e.g., wait, left, or right) to direct a bus qubit along a unique path to the target memory cell. This structure ensures that only O(log n) routing elements are activated for any given query, and, depending on the control model, the associated gate and latency overhead scales polylogarithmically (commonly reported between O(log n) and O(log^2 n)), rather than linearly with memory size. Such scaling mitigates the hardware fan-out that could otherwise negate quantum speedups (19). Robustness studies further indicate that errors in the bucket-brigade tend to remain localized along the activated path, offering insight into noise propagation and informing requirements for fault-tolerant implementation (17). In the context of our work, this capability is critical. Our algorithms load the DNA substring beginning at position I for all i in superposition, and the availability of an efficient QRAM, such as one implemented via bucket-brigade routing, is essential to avoid data-access bottlenecks. Accordingly, our complexity analysis assumes the existence of such an idealized, high-performance QRAM.

An ideal QRAM can, in principle take an address superposition \(\sum_i \alpha_i \ket{i}\) and return \[\sum_i \alpha_i \ket{i}\ket{\text{data}[i]}\] in O(log n) or even O(1) time per query (depending on the model). This means that the substring for any address can be loaded on demand during the oracle evaluation, without requiring all possibilities to be precomputed.

### 2.3 Quantum computing framework and implementation

All quantum circuits were implemented using Qiskit (version 0.45+) (23) with the Aer quantum simulator backend which provides a robust platform for developing and testing quantum algorithms before deployment on actual quantum hardware. The circuits employ standard quantum gates including Hadamard gates (H) for creating superposition states, Pauli-X gates for bit flips, controlled-NOT gates (CNOT) for entanglement operations, and multi-controlled-X gates (MCX) for implementing complex conditional logic operations that form the backbone of the quantum oracles (Figure 3A, 3B).

DNA sequences were encoded using a 2-bit quantum representation where each nucleotide maps to a unique binary pattern. Adenine maps to (0,0), Cytosine to (0,1), Guanine to (1,0), and Thymine to (1,1). This encoding scheme allows a k-mer of length L to be represented using exactly 2L qubits, enabling efficient quantum manipulation of genetic sequences while maintaining the full information content of the original DNA data. The choice of this particular encoding ensured that all nucleotides are equally represented in the quantum state space and facilitated straightforward quantum operations for sequence comparison.

Both algorithms utilize QRAM for storing and querying DNA sequence data, representing one of the most sophisticated aspects of the implementation. However, they differ in their pattern storage approaches. The *Enumerate-m* algorithm uses direct pattern encoding within the quantum oracle, while the *Nested Grover* algorithm employs a separate QRAM system for pattern storage. QRAM enables quantum superposition over memory addresses, allowing simultaneous access to multiple sequence positions, a capability that forms the foundation for the quantum speedup achieved by both algorithms. The QRAM architecture stores text substrings of length L at S = n - L + 1 addressable positions, where n represents the total sequence length, ensuring that every possible k-mer position in the target sequence can be accessed through the quantum addressing system.

### 2.4 The *Enumerate-m* algorithm: sequential pattern matching in quantum superposition

The *Enumerate-m* algorithm (Figure 1A, 1B) represents the more intuitive of the two quantum approaches, extending classical pattern matching concepts into the quantum domain by simultaneously checking a data substring against all m target patterns. This algorithm builds upon the foundational principles of Grover’s search algorithm but adapts them to handle multiple pattern matching through sequential enumeration within each quantum iteration. The theoretical foundation rests on the insight that while quantum superposition allows simultaneous access to all possible substring positions, the pattern matching process can still follow a classical enumeration strategy within the quantum oracle.

**Figure 1A.**
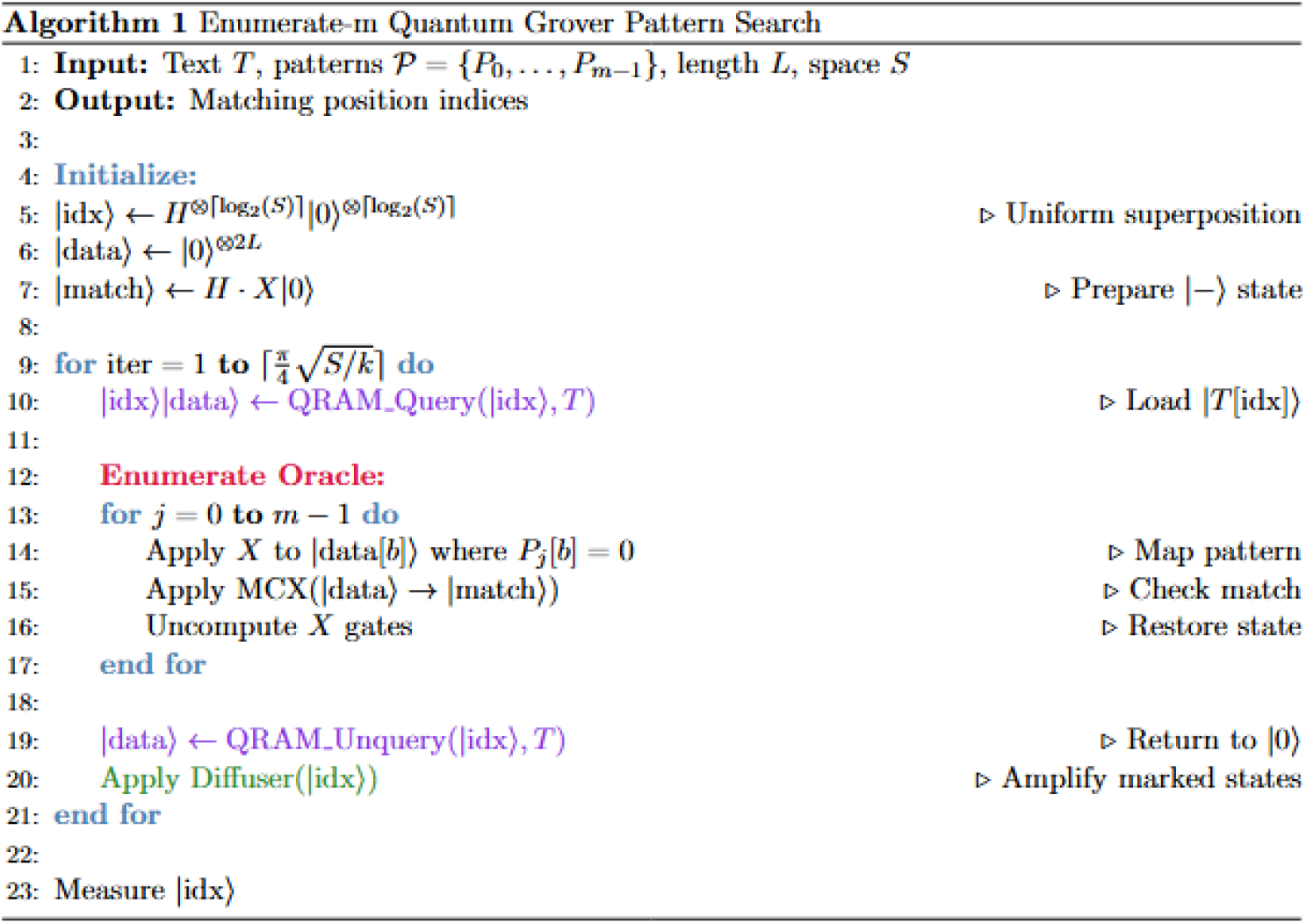
**A.** Algorithmic overview of the *Enumerate*-*m* algorithm

**Figure 1B.**
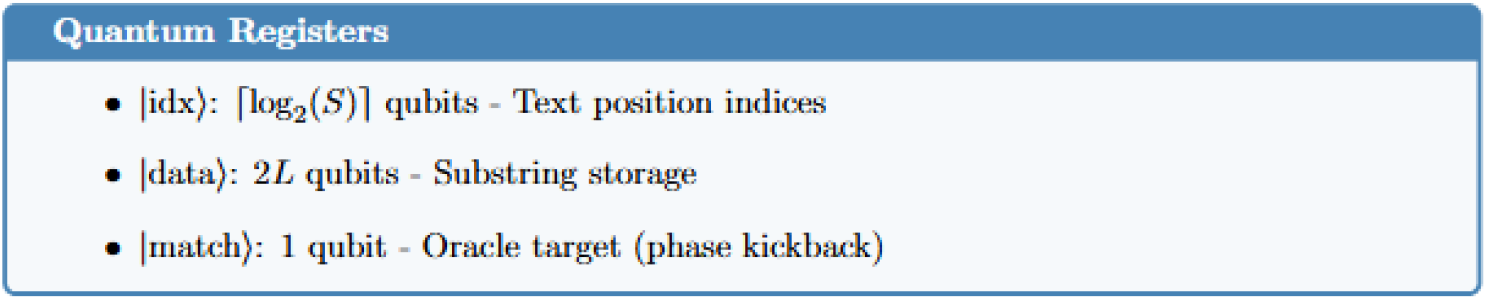
Quantum register requirements of the *Enumerate*-*m* algorithm

The quantum circuit architecture for the Enumerate-m algorithm consists of several carefully designed quantum registers that work in concert to achieve the desired functionality. The address register, requiring [log_2_(S)] qubits, encodes all possible substring positions within the target sequence, allowing the algorithm to simultaneously consider every location where a k-mer might occur. The data register uses 2L qubits to store the current substring loaded from the quantum memory, maintaining the full nucleotide information in the quantum 2-bit encoding scheme. A single-qubit match register indicates whether any of the target patterns have been found at the current superposed positions, serving as the output mechanism for the quantum oracle. Additional ancilla qubits support the complex multi-controlled operations required for efficient quantum computation, with their number determined by the largest multi-controlled operation in the circuit.

The algorithm proceeds through a carefully orchestrated sequence of quantum operations that begins with initialization of the address register in a uniform superposition over all S possible substring positions, creating the quantum state |ψ⟩ = (1/√S) ∑_i=0_^{S-1} |i⟩. This superposition represents the quantum advantage, as all possible starting positions for k-mer matches are explored simultaneously rather than sequentially as in classical algorithms. The match register is prepared in the |−⟩ state (|0⟩ - |1⟩)/√2 to enable phase kickback during oracle operations, a standard technique in quantum algorithms that ensures amplitude amplification affects the correct quantum states.

Each iteration of the *Enumerate-m* algorithm follows a compute-oracle-uncompute pattern that maintains quantum coherence while performing the necessary pattern matching operations. The QRAM query operation loads the substring at each superposed address into the data register, creating an entangled state between positions and their corresponding sequence data. The pattern oracle then sequentially tests the loaded substring against each of the m target patterns using a sophisticated quantum comparison mechanism. For each pattern p, the oracle applies Pauli-X gates to data qubits corresponding to ‘0’ bits in the pattern, effectively creating a quantum state where pattern matching corresponds to having all data qubits in the |1⟩ state. A multi-controlled-X gate with all data qubits as controls and the match qubit as target performs the actual matching test, marking the desired state. This is followed by uncomputation of the Pauli-X operations, which reverses the earlier transformations and restores the data qubits to their original state, ensuring the circuit remains reversible and free of residual entanglement.

The quantum diffusion operator (Figure 2C), implementing the inversion-about-average operation central to Grover’s algorithm, amplifies the amplitudes of states corresponding to matching addresses while diminishing those of non-matching states. After O(√S) iterations of this process, the address register is measured to obtain the positions where k-mer matches are most likely to occur. The algorithmic complexity reveals both the strengths and limitations of this approach. While the oracle complexity per iteration scales as O(m·L) due to sequential pattern checking, the total quantum gate count of O(√S · m · L) for amplitude amplification still provides a quadratic improvement over classical approaches in the sequence length dimension.

**Figure 2A.**
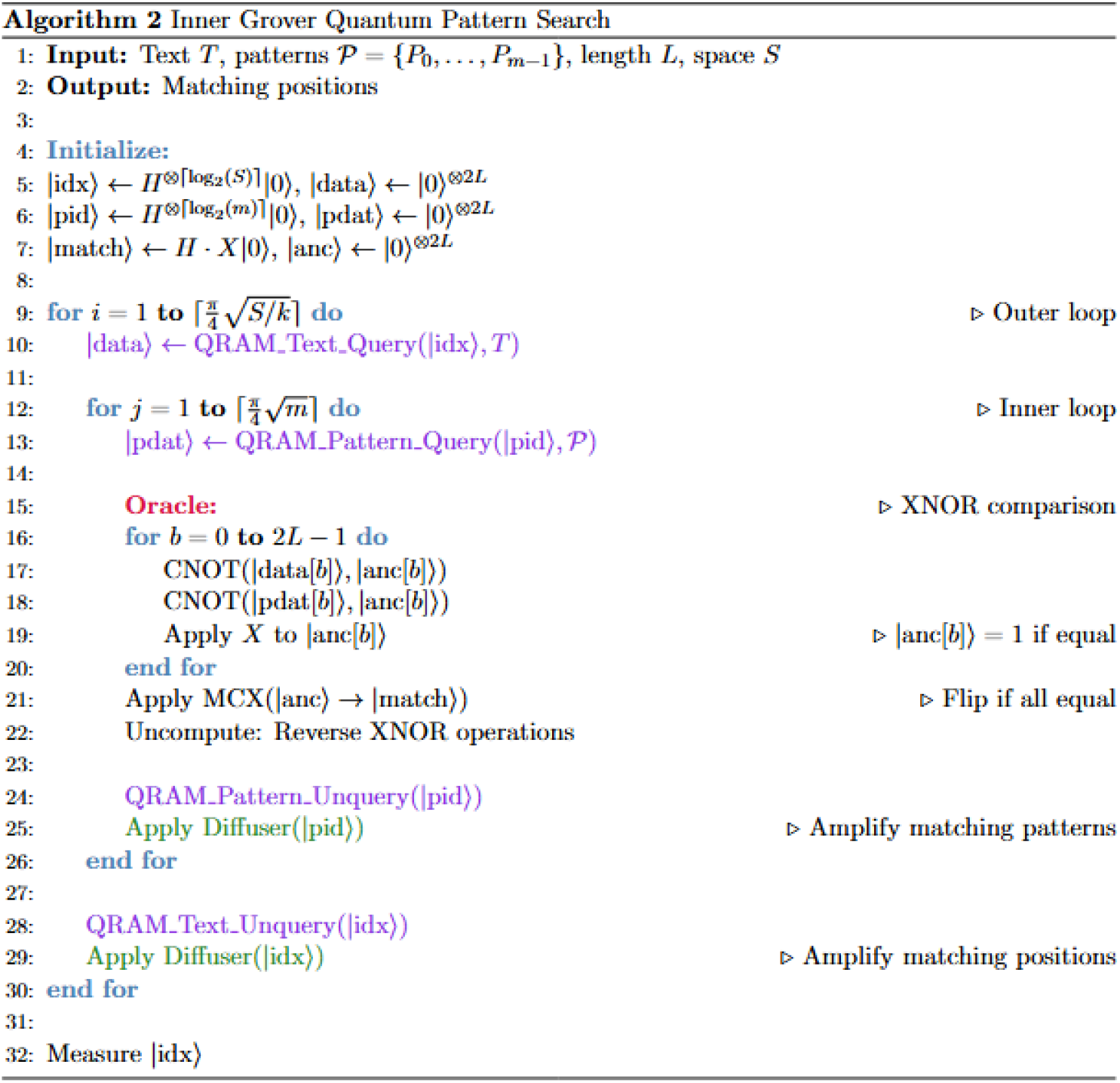
**A.** Algorithmic outline of the *Nested Grover* algorithm

**Figure 2B.**
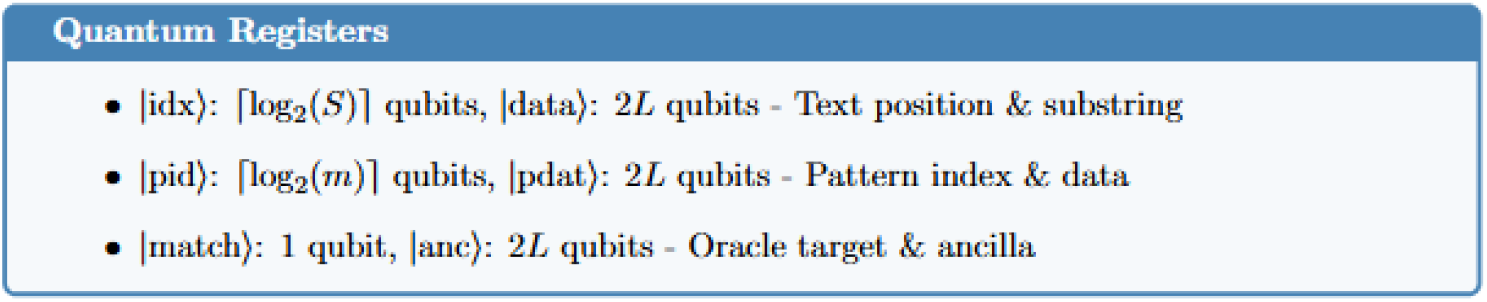
**B.** Quantum register requirements of the *Nested Grover* algorithm

**Figure 2C.**
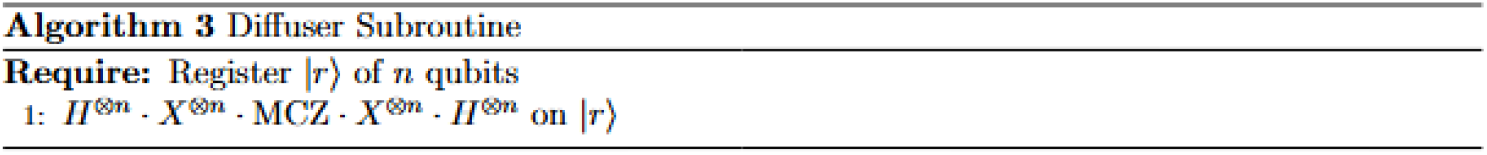
**C.** The diffuser subroutine is identical in both algorithms

### 2.5 The *Nested Grover* algorithm: layered quantum search architecture

The *Nested Grover* algorithm (Figure 2A, 2B) represents a more sophisticated approach that employs a nested amplitude amplification strategy, combining an inner Grover search over the pattern space with an outer Grover search over sequence positions. This hierarchical design builds on the insight that both the sequence position space and the pattern space can benefit from quantum parallelism, leading to a two-dimensional quantum search that can achieve superior scaling properties for large pattern databases. The theoretical foundation is grounded in the principle that nested applications of Grover’s algorithm can provide multiplicative improvements when searching through multiple dimensions simultaneously.

The circuit architecture extends beyond the *Enumerate-m* design by introducing additional quantum registers that enable the dual search capability. The pattern ID register, requiring ⌈log_2_(m)⌉ qubits, encodes indices for all target patterns, allowing the algorithm to simultaneously consider every pattern in the database. A pattern data register using 2L qubits stores the current pattern loaded from the pattern database, working in parallel with the sequence data register to enable direct quantum comparison between loaded sequence segments and search motifs. Specialized equality ancilla qubits, numbering 2L, support the explicit pattern comparison operations that form the core of the inner oracle. The total qubit requirement significantly exceeds that of the *Enumerate-m* algorithm, reflecting the increased computational power needed to achieve the better asymptotic scaling (Figure 4B).

The algorithm begins with a more complex initialization phase that prepares both the address and pattern ID registers in uniform superposition, creating the quantum state |ψ⟩ = (1/√(S·m)) ∑_i=0_^{S-1} ∑ □_i=0_^{m-1} |i⟩|j⟩. This initialization establishes superposition over all possible combinations of sequence positions and patterns, setting the stage for the nested search structure. The match register is again prepared in the |−⟩ state to enable proper phase kickback during oracle operations.

The layered loop structure distinguishes the *Nested Grover* algorithm from its simpler counterpart through a sophisticated interplay between outer and inner quantum searches. The outer loop, running for O(√S) iterations, focuses on amplitude amplification over sequence positions. Within each outer iteration, the algorithm first loads the substring at all superposed address positions using the QRAM text query operation. The inner loop then performs approximately ⌈π√m/4⌉ iterations to amplify the amplitudes of patterns that match the loaded substring. Each inner iteration involves loading a pattern at superposed pattern indices, comparing the loaded substring with the loaded pattern using an explicit equality oracle, unloading the pattern data to maintain phase-only oracle behavior, and applying a Grover diffusion operation over the pattern ID register (Figure 2C).

The equality oracle implementation represents one of the most technically sophisticated aspects of the *Nested Grover* algorithm, performing quantum comparison between the data and pattern registers through a carefully designed sequence of operations. The oracle constructs XNOR bits for each position by applying CNOT gates from both data and pattern qubits to dedicated ancilla qubits, then inverting these ancillas to convert XOR operations into XNOR operations that output |1⟩ when the corresponding bits are equal. A multi-controlled-X gate with all equality ancilla qubits as controls and the match qubit as target then performs the final comparison, followed by uncomputation of the XNOR operations to clean the ancilla qubits for subsequent use.

After completing the inner search, the algorithm unloads the substring data and applies an outer Grover diffusion operation over the address register to amplify addresses where pattern matches were found. The complexity analysis reveals the algorithm’s key advantage: an oracle complexity of O(L·√m) per outer iteration, leading to a total quantum gate count of O(√S · L · √m) for the complete nested amplitude amplification process. This represents a significant improvement over the *Enumerate-m* algorithm when the number of patterns m becomes large, as the √m scaling replaces the linear m scaling in the oracle complexity.

### 2.6 Comparative analysis of the two quantum approaches

The fundamental differences between the *Enumerate-m* and *Nester Grover* algorithms stem from their contrasting philosophies regarding how to exploit quantum parallelism in multi-pattern search problems. The *Enumerate-m* algorithm maintains the classical paradigm of sequential pattern checking while leveraging quantum superposition only in the sequence position dimension. In contrast, the *Nested Grover* approach fully embraces quantum parallelism by creating superposition states over both sequence positions and pattern indices, enabling simultaneous exploration of both dimensions through nested amplitude amplification.

The algorithmic complexity comparison reveals the critical trade-off between simplicity and asymptotic efficiency. The *Enumerate-m* algorithm processes patterns sequentially within each quantum iteration, resulting in oracle complexity that scales linearly as O(m·L) per iteration, where m represents the number of patterns and L the k-mer length. This linear scaling with pattern count means that the algorithm’s efficiency degrades proportionally as the pattern database grows, ultimately limiting its applicability to relatively small pattern sets. The *Nested Grover* algorithm, through its multi-level search structure, achieves oracle complexity of O(L·√m) per outer iteration, representing a fundamental improvement for large pattern databases where the square-root scaling provides substantial advantages over linear enumeration.

Circuit complexity considerations reveal additional dimensions of the trade-off between the two approaches (Figure 3A, 3B). The *Enumerate-m* algorithm requires a relatively modest number of qubits: 2L qubits for data storage, ⌈log_2_(S)⌉ qubits for address encoding, and additional ancilla qubits for multi-controlled operations. The circuit depth remains moderate due to the straightforward sequential structure of pattern checking operations. The Inner Grover algorithm demands significantly more quantum resources, requiring an additional ⌈log_2_(m)⌉ qubits for pattern indexing, 2L qubits for pattern data storage, and 2L dedicated ancilla qubits for equality checking operations. The nested loop structure also increases circuit depth considerably, potentially making the algorithm more susceptible to decoherence and gate errors in near-term quantum devices.

**Figure 3.**
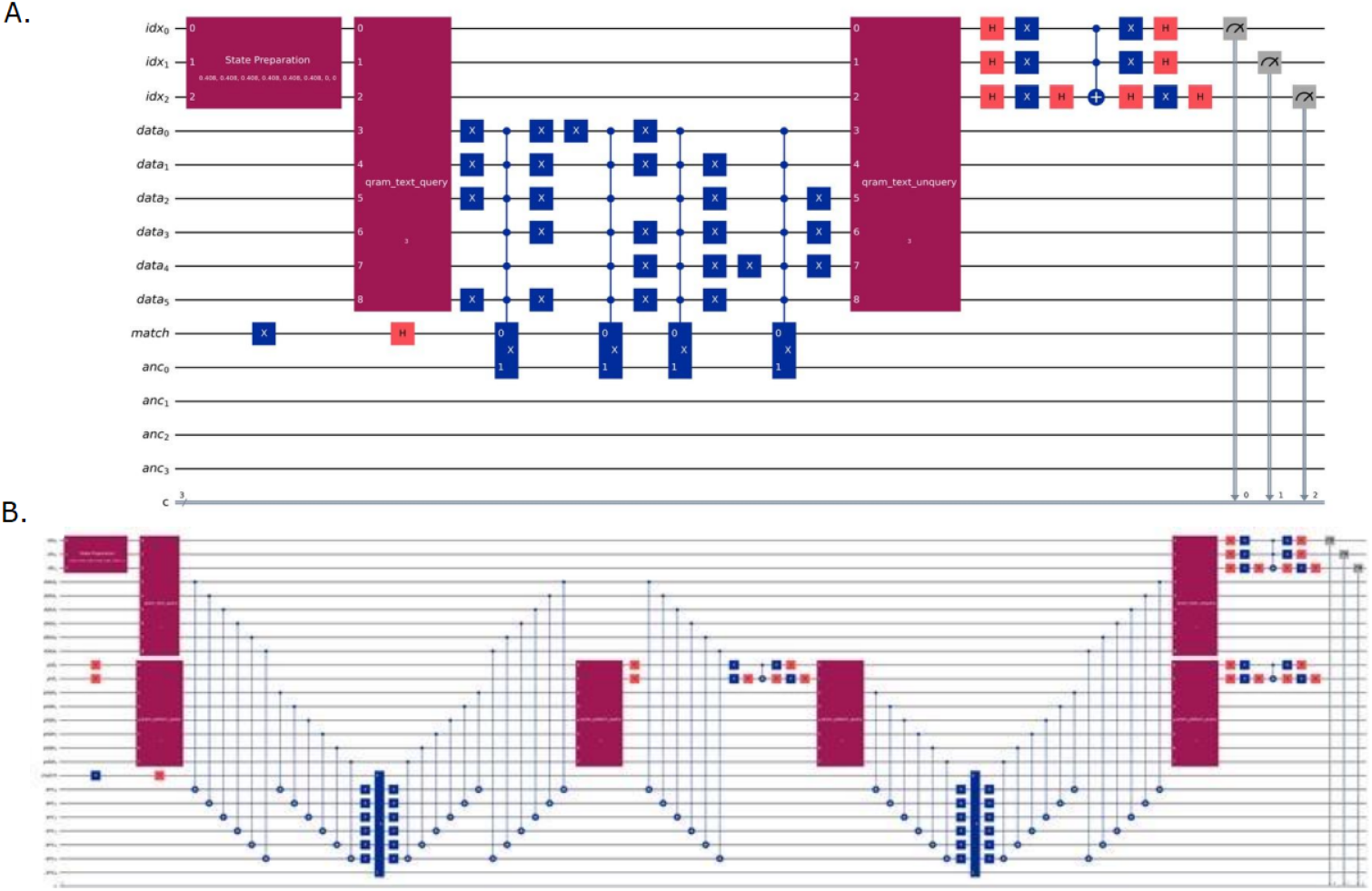
**A.** Quantum circuit representations of *Enumerate*-*m* under the ideal QRAM assumption. **B**. Quantum circuit representations of *Nested Grover* under the ideal QRAM assumption.

The practical implications of these differences become particularly pronounced when considering real-world applications. For genomic applications involving small to moderate numbers of k-mer patterns, such as searching for specific regulatory sequences or known mutations, the *Enumerate-m* algorithm offers a compelling balance of simplicity and performance. Its lower resource requirements and straightforward implementation make it more suitable for near-term quantum devices with limited qubit counts and high error rates. The algorithm’s resilience to noise stems from its simpler structure and shorter circuit depth, reducing the cumulative impact of gate errors and decoherence.

For large-scale genomic analyses involving extensive pattern databases, such as comprehensive k-mer frequency analysis or whole-genome pattern searches, the *Nested Grover* algorithm’s superior asymptotic scaling becomes increasingly important. Despite its higher resource requirements and increased complexity, the square-root improvement in pattern processing can provide substantial speedups when dealing with thousands or millions of patterns. However, the practical realization of these theoretical advantages depends critically on the availability of fault-tolerant quantum computers capable of supporting the large number of qubits and deep circuits required by the nested algorithm structure.

The ancilla management requirements represent another crucial practical consideration distinguishing the two approaches. The *Enumerate-m* algorithm’s ancilla needs scale primarily with the largest multi-controlled operation in the circuit, typically requiring ancillas proportional to the maximum of the address register size and data register size. The *Nested Grover* algorithm’s ancilla requirements are more substantial and multifaceted, needing 2L dedicated ancillas for equality checking operations plus additional ancillas for decomposing the various multi-controlled operations throughout the nested structure. This increased ancilla overhead directly impacts the algorithm’s feasibility on resource-constrained quantum hardware.

Error propagation characteristics further differentiate the two algorithms in practical implementations. The *Enumerate-m* algorithm’s linear structure means that errors in individual gates affect the computation in relatively predictable ways, and the algorithm maintains some robustness to isolated gate failures. The layered structure of the *Nested Grover* algorithm creates more complex error propagation patterns, where errors in the inner loop can accumulate and amplify through multiple iterations of the outer loop. This amplification effect potentially requires more sophisticated error correction schemes to achieve reliable operation, adding another layer of complexity to practical implementations.

### 2.7 Theoretical speedup in genomic pattern matching

The classical Aho-Corasick algorithm offers optimal search performance with a time complexity of O(n + m), where n is the length of the input text and m is the total length of all search patterns. This linear runtime has long been the benchmark for multi-pattern string matching. In contrast, our quantum implementation, assuming ideal QRAM, introduces two variants with distinct complexity profiles. The *Enumerate-m* variant requires O(\sqrt{n}) Grover iterations, each costing O(L \cdot m), leading to a total gate complexity of O(\sqrt{n} \cdot L \cdot m). The *Nested Grover* variant, more efficient in pattern handling, performs O(\sqrt{n}) outer iterations with a per-iteration cost of O(L \cdot \sqrt{m}), resulting in a gate complexity of O(\sqrt{n} \cdot L \cdot \sqrt{m}).

A theoretical quantum advantage emerges when the gate complexity of the *Nested Grover* variant falls below the classical linear bound, specifically, when \sqrt{n} \cdot L \cdot \sqrt{m} < n. This condition is achievable in scenarios involving extremely large genomic datasets (e.g., n \approx 10^8) and a substantial number of patterns (e.g., m \approx 10^4), assuming motif lengths L remain moderate. Under these parameters, the quantum algorithm offers a potential quadratic speedup over classical methods. While current limitations, particularly QRAM overhead and gate depth, pose practical challenges, the analysis provides a rigorous foundation for future quantum-enhanced approaches to large-scale string matching.

### 2.8 QRAM and the path to scalable quantum search

QRAM offers a principled framework for mitigating the data-loading bottleneck in quantum search algorithms by enabling superposition-based access to classical memory. This capability allows substrings corresponding to all candidate indices to be retrieved in parallel, reducing the asymptotic data-access cost from O(S \cdot L) explicit gate operations to O(\log S) address qubits plus the physical cost of a single query. If realised on low-latency, fault-tolerant hardware, this approach could significantly reduce circuit depth and qubit overhead by enabling dynamic data access, thereby removing the need to store all patterns or substrings within the circuit. It would also allow parallel pattern matching across large search spaces without the linear increase in gate count typically associated with such tasks.

While QRAM is still in the process of being engineered for practical use, its theoretical appeal continues to inspire active research. Practical architectures are yet to be fully established, but ongoing progress in circuit-level optimizations, oracle depth reduction, and hybrid quantum–classical batching techniques is steadily moving the field forward. These developments are key to transforming proof-of-concept implementations into scalable quantum bioinformatics tools capable of addressing genome-scale datasets.

### 2.9 Implementation challenges and practical considerations

The transition from theoretical quantum algorithms to practical implementations reveals several critical challenges that significantly impact the feasibility and performance of both approaches on current and near-term quantum hardware. Multi-controlled-X gates, which form the backbone of both algorithms’ oracle implementations, present one of the most significant practical hurdles. When decomposed into elementary quantum gates available on physical quantum devices, an MCX gate with n controls requires O(n) elementary gates and potentially O(n) ancilla qubits for efficient implementation. This decomposition overhead can dramatically increase the actual circuit depth and resource requirements beyond the theoretical analysis, particularly affecting the *Nested Grover* algorithm with its more extensive use of large MCX operations.

The gate complexity implications extend beyond simple counting to fundamental questions about quantum circuit optimization and compilation. Current quantum compilers must balance the trade-off between minimizing circuit depth (to reduce decoherence effects) and minimizing gate count (to reduce error accumulation), often leading to suboptimal implementations of the high-level algorithmic structures. The QRAM operations, while conceptually elegant, require sophisticated implementations that may not be directly available on near-term quantum hardware, necessitating approximations or alternative approaches that could significantly alter the algorithms’ performance characteristics.

Connectivity constraints on physical quantum devices add another layer of complexity to the implementation challenge. Most current quantum processors have limited qubit connectivity, meaning that not all qubits can directly interact with all others. This limitation forces the insertion of additional SWAP gates to move quantum information between distant qubits, increasing both circuit depth and error rates. The *Nested Grover* algorithm, with its more complex register interactions and extensive ancilla requirements, is particularly vulnerable to connectivity limitations that could make efficient implementation difficult or impossible on current hardware architectures.

The noise characteristics of near-term quantum devices profoundly impact the practical viability of both algorithms. Gate errors, decoherence, and measurement errors accumulate throughout the quantum computation, with longer and deeper circuits suffering more severe degradation. The *Enumerate-m* algorithm’s shorter circuit depth and simpler structure provide some inherent resilience to these noise sources, potentially enabling successful operation even on relatively noisy quantum devices. The *Nested Grover* algorithm’s layered structure and longer coherence requirements may necessitate error correction schemes or highly optimized implementations to achieve practical utility.

Calibration and control challenges represent additional practical considerations that can significantly impact algorithm performance. Quantum devices require frequent recalibration to maintain gate fidelities and minimize crosstalk between qubits. The *Nested Grover* algorithm’s complex nested loop structure and precise timing requirements may be more sensitive to calibration drift and control errors, potentially requiring more frequent recalibration cycles or more sophisticated control protocols. The algorithm’s dependence on precise amplitude amplification through multiple nested iterations means that small errors in individual gates can compound to produce significant deviations from the intended quantum state evolution.

## 3. Results

To demonstrate the principles and performance of our quantum string matching implementations, we evaluated them on two trivial DNA pattern-matching tasks of low complexity. The simplest illustrative example involved searching three 2-mers, “AC”, “CG” and “GT”, within an 4-base DNA sequence “ACGT”, yielding three possible start positions (S = 3) and one outer Grover iteration (R = 1). DNA bases were encoded using two qubits each, A = (0,0), C = (0,1), G = (1,0), T = (1,1), so a pattern of length L required 2L qubits. This encoding was used to load substrings into quantum registers via QRAM-style operations. This small-scale use case allowed us to illustrate the full quantum circuit structures (Figure 3A, 3B), including initialization of superposition states, QRAM queries to load substrings and patterns, oracle marking of matching states, QRAM uncomputation to preserve phase-only oracles, and diffusion operators for amplitude amplification. It further enabled us to estimate the qubit requirements of each implementation and to gauge the complexity, even at a toy-scale level (Figure 4A, 4B).

**Figure 4.**
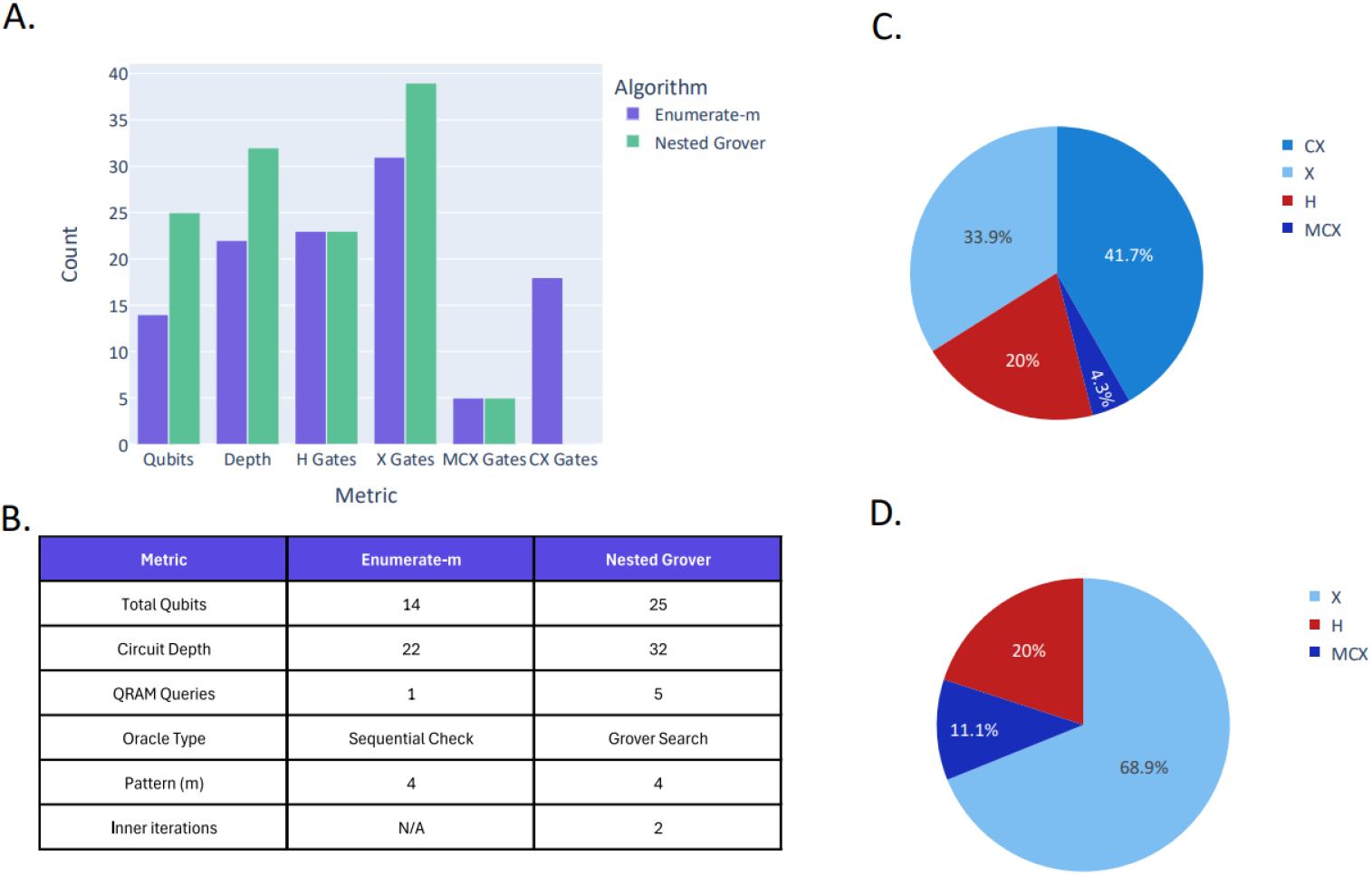
**A–B.** Circuit statistics comparison between the two proposed methods using the toy example text “ACGT” with patterns “AC”, “CG”, and “GT”. **C**. Gate distribution for *Enumerate*-*m*. **D**. Gate distribution for *Nested Grover*.

The two algorithmic implementations, *Enumerate-m* and *Nested Grover*, demonstrated distinct trade-offs. The *Enumerate-m* sequentially checked all m patterns against the loaded substring, requiring O(m·L) work per oracle call but maintaining a simpler circuit structure with fewer quantum registers (Figure 3A). The *Nested Grover* algorithm incorporated an inner Grover loop over the pattern space, achieving O(√m) speedup at the cost of additional qubit overhead for pattern index and pattern data registers (Figure 3B). Circuit statistics present clearly these architectural differences (Figure 4A, 4B, 4C, 4D). While the *Enumerate-m* approach used fewer total qubits, the *Nested Grover* method scaled more efficiently for large pattern sets due to its O(L) oracle complexity and dual amplitude amplification over both text and pattern spaces. Gate distribution analysis revealed multi-controlled X (MCX) gates as the dominant resource cost, since they appear repeatedly in oracle constructions, diffusion operators, and QRAM routines. Unlike single-or two-qubit gates, MCX gates require a large number of ancillary qubits and decompositions into many basic operations, making them far more expensive in depth and error accumulation. This overhead often represents the main barrier to scaling, as MCX gates are difficult to implement fault-tolerantly and tend to stress hardware connectivity (24). In the above simplified scenario (text “ACGT”, patterns “AC”, “CG”, “GT”), the circuits remained compact and suitable for near-term quantum devices (Figure 4B).

To further assess implementation performance, we conducted a second, moderately larger experiment designed to evaluate algorithmic robustness. The target sequence “ACGTACGT” served as the base text, with motifs “GTA” and “TAC” specified as substrings to be detected. As illustrated in Figure 5, the *Enumerate*-*m* implementation exhibited greater variability, yet produced index positions that were readily distinguishable from background noise. In contrast, the *Nested Grover* implementation yielded only the precise motif indices (starting positions). It merits attention that, the *Nested Grover* approach interprets the base text cyclically, overflowing into index 0 after the terminal (7th) position. Although it achieves a more coherent and consistently robust performance, this advantage is accompanied by a significant overhead in gate-level circuit complexity.

**Figure 5.**
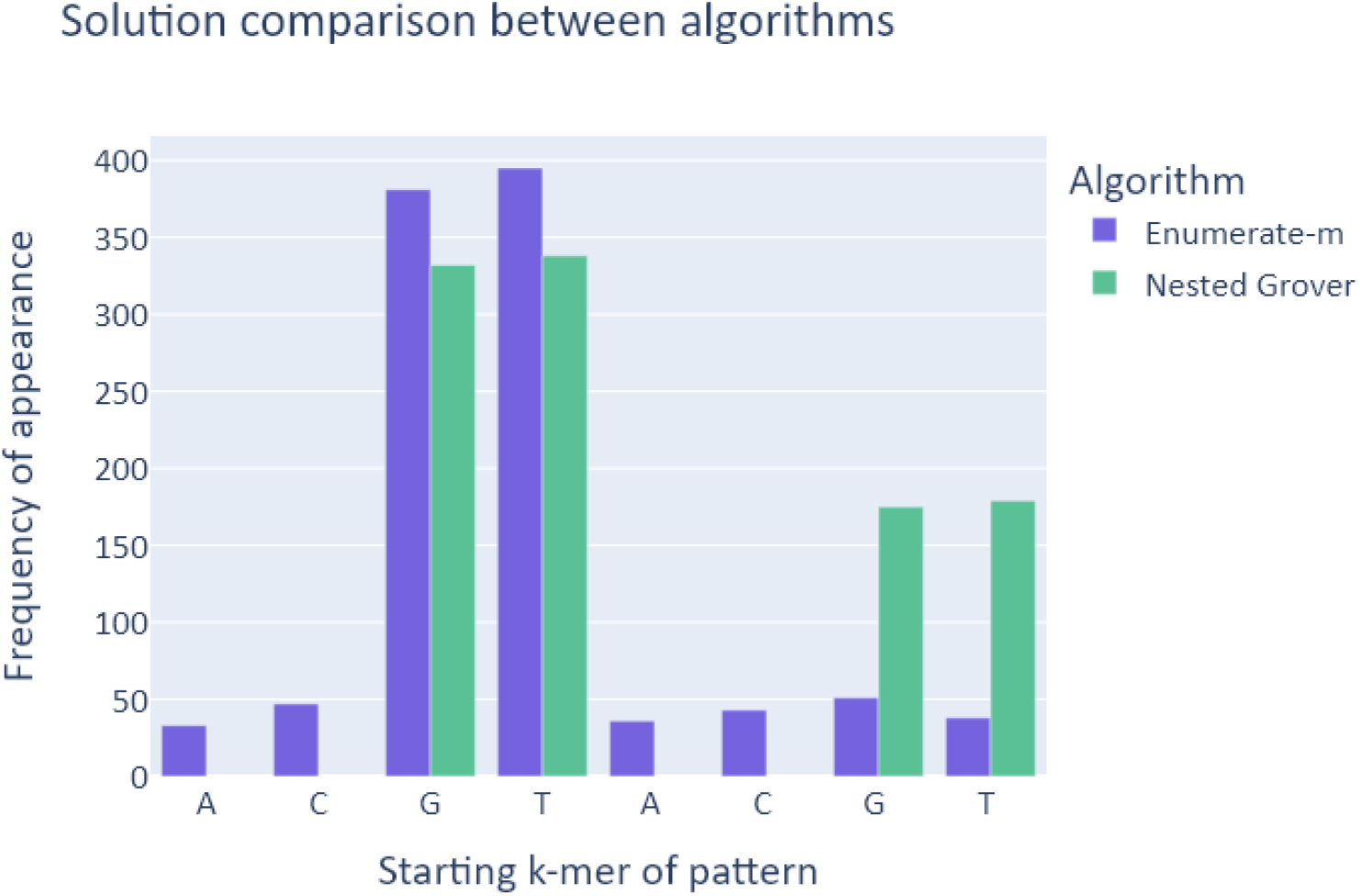
Comparison of the *Enumerate*-*m* and *Nested Grover* implementations on the target sequence “ACGTACGT” with motifs “GTA” and “TAC.” The X-axis represents positions within the base text (0 = A, 1 = C, 2 = G, etc.), whereas the Y-axis denotes the frequency with which the algorithm correctly detected the queried motifs’ starting position across 1,024 trials per algorithm.

To quantify scalability, we analyzed larger genomic workloads. Searching a 1000-base DNA sequence against 10 patterns of length 20 nucleotides requires an index register of 10 qubits (981 positions), a 40-qubit character register (2 qubits per nucleotide), and additional ancillas for phase inversion, match indication, and intermediate computation, totaling approximately 53 qubits per search instance. The oracle construction alone involves 19,620 MCX gates (981 positions × 20 bases), each decomposing into many elementary gates, plus hundreds of Toffoli and X gates per pattern. These figures underscore the steep resource demands even for modest genomic inputs.

From a more practical perspective, scaling to human chromosome 1 (∼250 million bases) with a pattern length of 20 would require an index register of at least 28 qubits to enumerate all S = 249,999,981 starting positions, plus 40 qubits for pattern encoding and ancillas, totaling around 72 qubits per search instance. Substring loading alone would demand on the order of 5 billion MCX gates, with total gate counts (including amplification and verification) reaching tens of billions. These depths far exceed the coherence times and gate fidelities of current Noisy Intermediate-Scale Quantum (NISQ) hardware, highlighting the need for error correction, gate-efficient oracle design, and hardware-aware compilation strategies to make such large-scale applications feasible.

The theoretical quantum advantage of these approaches stems from parallel evaluation of all text positions through superposition and Grover’s amplitude amplification, achieving O(√S) query complexity compared to classical O(S) exhaustive search. The *Nested Grover* algorithm extends this advantage by applying Grover search to the pattern space as well, yielding combined O(√S · √m) iteration complexity with O(L) work per oracle call, compared to *Enumerate-m*’s O(√S) iterations with O(m·L) oracle cost. However, these speedups critically depend on efficient QRAM implementation, which remains an open research problem in quantum computing. Without practical QRAM solutions, the explicit gate expansion shown in the demo circuits reveals substantial overhead that may diminish or eliminate the theoretical quantum advantage, particularly for the *Nested Grover* approach with its higher gate count and circuit depth.

## 4 Discussion

This study introduces two novel quantum algorithms for multi-pattern string matching that demonstrate theoretical performance improvements over classical approaches, including the widely used Aho-Corasick algorithm. By leveraging quantum principles such as superposition and amplitude amplification, these algorithms offer a fundamentally different computational paradigm that holds promise for accelerating search tasks in high-dimensional biological datasets.

Our simulations demonstrate that quantum search primitives, particularly those inspired by Grover’s algorithm, can reduce search complexity under idealized conditions. Realizing these gains in practice depends on efficient quantum memory access, as QRAM operations introduce additional overhead. Current hardware is still evolving toward the scale and fidelity needed for realistic genomic workloads, but ongoing improvements in architecture and error mitigation are steadily advancing the feasibility of such applications. Rather than representing a limitation, these observations underscore the importance of hardware-aware algorithmic design and point to specific scenarios, such as extremely large pattern sets or highly repetitive queries, where quantum advantage may be achievable in the near term.

Hybrid quantum-classical strategies emerge as a promising direction, particularly in the context of NISQ devices. These approaches allow partial acceleration through shallow quantum circuits and modular problem decomposition, while retaining the robustness of classical methods. Such strategies may offer practical benefits without requiring full fault-tolerant quantum computing.

The successful integration of quantum algorithms into biological workflows will require close collaboration with domain experts. Understanding the biological relevance of motif sizes, sequence structures, and search tolerances is essential for tailoring algorithmic solutions to real-world applications. Furthermore, evaluating performance using biologically meaningful metrics, rather than relying solely on raw computational speed, will be essential for demonstrating the true value of quantum algorithms. This approach ensures that their insights are not only computationally impressive but also practically actionable and impactful for advancing biological research.

Overall, this work adds to the development of a new class of bioinformatics tools. It invites further research into scalable quantum memory architectures, biologically informed algorithm design, and robust simulation frameworks. As quantum technologies continue to evolve, so too will the opportunities to interrogate genomic data with unprecedented efficiency and precision. Taken together, these advances signal that we are approaching the era of quantum bioinformatics, a paradigm in which quantum computing principles are seamlessly integrated into biological data analysis, enabling discoveries at scales and speeds previously unattainable.

## Code Availability

The source code supporting this work, along with detailed instructions on how to install dependencies and run the provided scripts, is openly available in the following GitHub repository [https://github.com/Georgakopoulos-Soares-lab/quantum-multi-motif-finder]. Users can access the repository to reproduce the results, adapt the workflows, and explore further applications

## Funding

Research reported in this publication was supported by the National Institute of General Medical Sciences of the National Institutes of Health under Award Number R35GM155468 and start-up funds to I.G.S.. The content is solely the responsibility of the authors and does not necessarily represent the official views of the National Institutes of Health.

**Author contributions**

Christos Papalitsas

(Conceptualization, Investigation, Methodology, Software, Visualization, Writing-original draft) Ioannis Mouratidis

(Conceptualization, Investigation, Writing-original draft)

Michail Patsakis

(Investigation, Writing-original draft)

Evangelos Stogiannos (Investigation, Visualization, Writing-original draft)

Ilias Georgakopoulos-Soares

(Conceptualization, Funding acquisition, Project administration, Writing-original draft) Grigorios Koulouras

(Conceptualization, Investigation, Project administration, Writing-original draft)

